# Tau filaments from multiple cases of sporadic and inherited Alzheimer’s disease adopt a common fold

**DOI:** 10.1101/411215

**Authors:** Benjamin Falcon, Wenjuan Zhang, Manuel Schweighauser, Alexey G. Murzin, Ruben Vidal, Holly J. Garringer, Bernardino Ghetti, Sjors H.W. Scheres, Michel Goedert

**Affiliations:** MRC Laboratory of Molecular Biology, Cambridge CB2 0QH, UK; Department of Pathology and Laboratory Medicine, Indiana University School of Medicine, Indianapolis, IN 46202

## Abstract

The ordered assembly of tau protein into abnormal filaments is a defining characteristic of Alzheimer’s disease (AD) and other neurodegenerative disorders. It is not known if the structures of tau filaments vary within, or between, the brains of individuals with AD. We used a combination of electron cryo-microscopy (cryo-EM) and immuno-gold negative-stain electron microscopy (immuno-EM) to determine the structures of paired helical filaments (PHFs) and straight filaments (SFs) from the frontal cortex of 17 cases of typical AD (15 sporadic and 2 inherited) and 2 cases of atypical AD (posterior cortical atrophy). The high-resolution structures of PHFs and SFs from the frontal cortex of 3 cases of typical AD, 2 sporadic and 1 inherited, were determined by cryo-EM. We also used immuno-EM to study the PHFs and SFs from a number of cortical and subcortical brain regions. PHFs outnumbered SFs in all AD cases. By cryo-EM, PHFs and SFs were made of two C-shaped protofilaments with a combined cross-β/β-helix structure, as described previously for a case of AD. The higher resolution structures obtained here showed two additional amino acids at each end of the protofilament structure. The immuno-EM findings, which established the presence of repeats 3 and 4, but not repeats 1 and 2, of tau in the filament cores of all AD cases and brain regions thereof, were consistent with the cryo-EM results. These findings show that there is no significant variation in tau filament structures between individuals with AD. This knowledge will be crucial for understanding the mechanisms that underlie tau filament formation and for developing novel diagnostics and therapies.

## INTRODUCTION

Alzheimer’s disease (AD) is the most common neurodegenerative disorder (1). The vast majority of cases are sporadic, with age being the main predisposing factor. Inheritance of the ε4 allele of the apolipoprotein E gene (*APOE*) is a major genetic risk factor for sporadic AD (2). Dominantly-inherited forms of AD, which are caused by mutations in the amyloid precursor protein gene (*APP*) (3,4) and the presenilin genes (*PSEN1* and *PSEN2*) (5,6) account for less than 1% of cases. No mechanism-based therapies exist.

All cases of AD are characterised by abundant intraneuronal neurofibrillary lesions of filamentous tau (7-10) and extracellular deposits of filamentous β-amyloid (Aβ) (11,12) in the cerebral cortex and other brain regions. Atypical forms of AD differ from typical forms by clinical presentation and the brain areas most severely affected by neurofibrillary lesions and Aβ deposits (13). Despite being made of unrelated proteins, tau and Aβ filaments share a cross-β architecture characteristic of amyloids (11,14). Filamentous tau inclusions, in the absence of Aβ deposits, define a number of other neurodegenerative diseases (15). Six tau isoforms are expressed in normal adult human brain: three isoforms have four microtubule-binding repeats (R1, R2, R3, R4; 4R tau) and three isoforms lack the second repeat (3R tau). Tau filaments of AD are composed of all six isoforms (16).

Neurofibrillary lesions consist of tangles in cell bodies, neuropil threads in the processes of nerve cells and dystrophic neurites associated with plaques. The accumulation of neurofibrillary lesions in brain follows a stereotypical pattern that correlates with atrophy and cognitive deficits (17). It follows that an understanding of the ordered assembly of tau into filaments is essential for diagnosis and therapy. Tau filaments from AD brain adopt two characteristic morphologies. Paired helical filaments (PHFs) constitute the major species; they have a helical crossover distance of approximately 70 nm, with pronounced variations in projected widths from 7 to 15 nm. Straight filaments (SFs), which have a similar crossover distance, but a constant width of approximately 10 nm, are in the minority (18).

The structures of PHFs and SFs extracted from the frontal cortex of an individual with sporadic AD were previously determined using electron cryo-microscopy (cryo-EM) (19). PHFs and SFs were found to be ultrastructural polymorphs, which share two identical protofilaments, but differ in their inter-protofilament packing. Each C-shaped protofilament adopted a combined cross-β/β-helix structure comprising residues 306-378 of human tau (in the numbering of the 441 residue isoform), i.e. R3, R4 and 10 amino acids following R4. The rest of the sequence was disordered and formed the fuzzy coat (10). Cryo-EM was also used to determine the structures of tau filaments from the fronto-temporal cortex of an individual with Pick’s disease (PiD) (20), a frontotemporal dementia with tau filaments made of 3R tau. The ordered core of tau filaments from PiD was made of a single protofilament with an elongated cross-β structure, comprising residues K254-F378 of tau (but lacking V275-S305 of R2). Distinct conformers of aggregated tau thus exist in human tauopathies.

It is not known if the structures of tau filaments vary within, or between, the brains of individuals with AD. This knowledge is vital for the development of therapies targeting tau assembly and the design of diagnostic ligands. It is also essential for understanding the mechanisms underlying tau filament formation and propagation.

Here, we used cryo-EM to determine the structures of tau filaments from two additional cases of sporadic AD and one case of dominantly-inherited disease (V717F mutation in *APP*). Since cryo-EM is too low-throughput to study a large number of cases, we also used immuno-gold negative-stain electron microscopy (immuno-EM) to determine the morphologies and tau repeat composition of the cores of PHFs and SFs from frontal cortex of the four cases used for cryo-EM; an additional twelve cases with the typical form of sporadic AD; and another case with dominantly-inherited disease. We also used immuno-EM to study tau filaments from temporal, occipital and cingulate cortices, as well as from thalamus, substantia innominata and putamen of AD case 1 used previously for cryo-EM (19); and for tau filaments extracted from frontal and occipital cortices of two cases of posterior cortical atrophy (PCA). PCA is an atypical form of sporadic AD with early visual dysfunction and neurodegeneration of posterior cortical regions (21,22). Our immuno-EM results show that the cores of PHFs and SFs from all AD cases and brain regions examined comprise R3 and R4 of tau, while our cryo-EM structures demonstrate that PHFs and SFs from frontal cortex of the three new cases of AD are like those we previously reported (19), indicating the existence of a common tau fold in AD.

## RESULTS

### Electron cryo-microscopy

We first used cryo-EM to image tau filaments extracted from the frontal cortex of an individual with sporadic AD (Figure 1, case 2 in Table 1). PHFs and SFs were present in a ratio of approximately 4:1. Using helical reconstruction in RELION (23), we determined the structures of PHFs to 3.2 Å and of SFs to 3.3 Å resolution (Figure 1, Online Resources 1, 2 and 3). As reported before (19) for filaments extracted from a case of sporadic AD (case 1 in Table 1), the core structures comprised V306-F378 of tau in a combined cross-β/β-helix, C-shaped fold (Online Resource 4). Moreover, the higher resolution structures of tau filaments reported here showed additional densities corresponding to G304 and S305 from R2 (or G273 and K274 from R1), as well as R379 and E380 from the sequence after R4, which were not as well-resolved in the structures from AD case 1 (19). The first β-strand of the protofilament structure begins at K274 of R1 or at S305 of R2 and ends at K311, whereas the eighth β-strand extends from N368-E380. Weaker densities extending from the N- and C- terminal regions of the core, described for filaments extracted from AD case 1 (19), were also observed, as were the densities interacting with the sidechains of K317, T319 and K321. Furthermore, in the protofilament interface of the PHF, the higher resolution structure revealed the presence of extra densities linking the side chains of K331 on one protofilament with the backbone atoms of V337 on the other protofilament (Online Resource 5). These densities may correspond to water molecules or a post-translational modification of K331, such as mono-methylation.

**Table 1:**
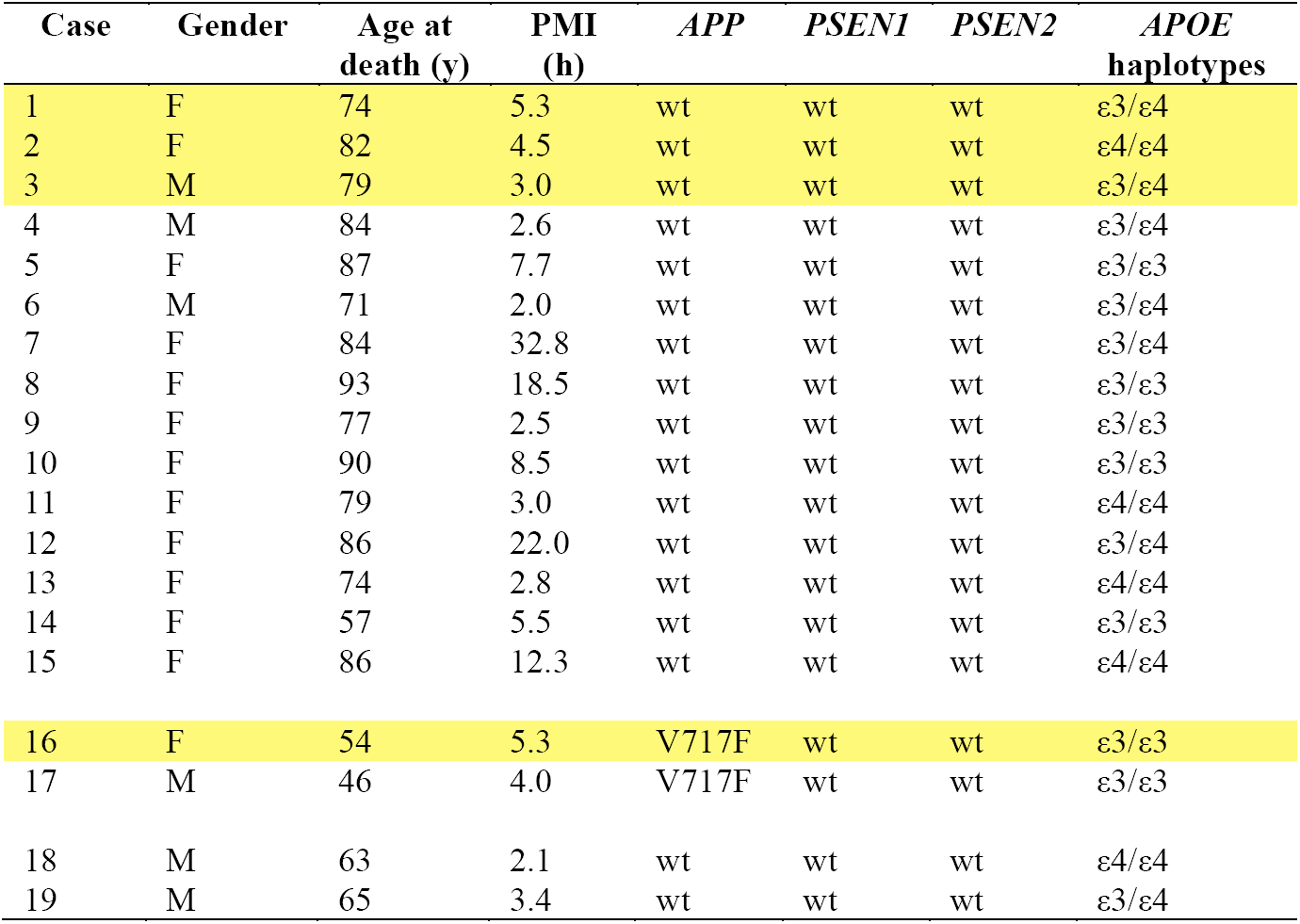
Cases of Alzheimer’s disease. Wild-type (wt), no known pathogenic mutations were detected. Cases 1, 2, 3 and 16 were used for cryo-EM (highlighted in yellow). PMI, post mortem interval.

**Figure 1:**
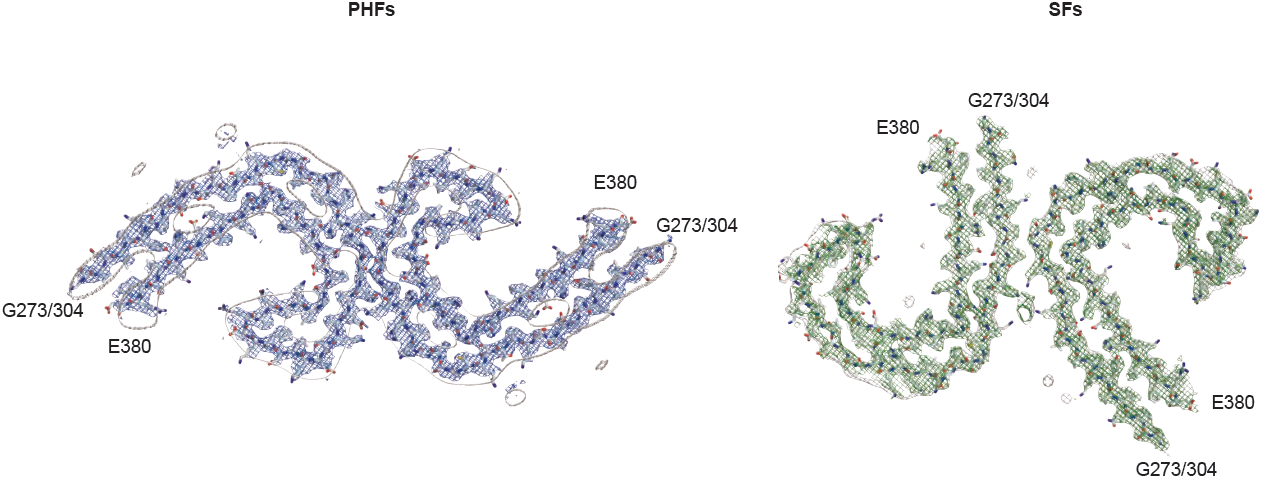
Cryo-EM densities and atomic models of PHFs and SFs from the frontal cortex of AD case 2. PHFs and SFs from case 2 are resolved to 3.2 Å and 3.3 Å, respectively. Sharpened, high-resolution maps are shown in blue (PHFs) and green (SFs). Unsharpened 4.5 Å low-pass filtered densities are shown in grey. The models comprise G273-E380 of 3R tau and G304- E380 of 4R tau.

We also used cryo-EM to determine the structures of tau filaments extracted from the frontal cortex of a third individual with sporadic AD (case 3 in Table 1) and an individual with dominantly-inherited AD (mutation V717F in *APP*, case 16 in Table 1). In both individuals, PHFs and SFs were present in a ratio of approximately 4:1. Despite lower resolution, the structures of PHFs and SFs were the same as those of tau filaments from cases 1 and 2 (Figure 2, Online Resource 2). A side-by-side comparison of PHF and SF reconstructions from cases 1-4 showed the common presence of extra densities bordering the solvent-exposed side chains of R349 and K375, and of H362 and K369 within each C-shape (Figure 2, Online Resource 2).

**Fig. 2.**
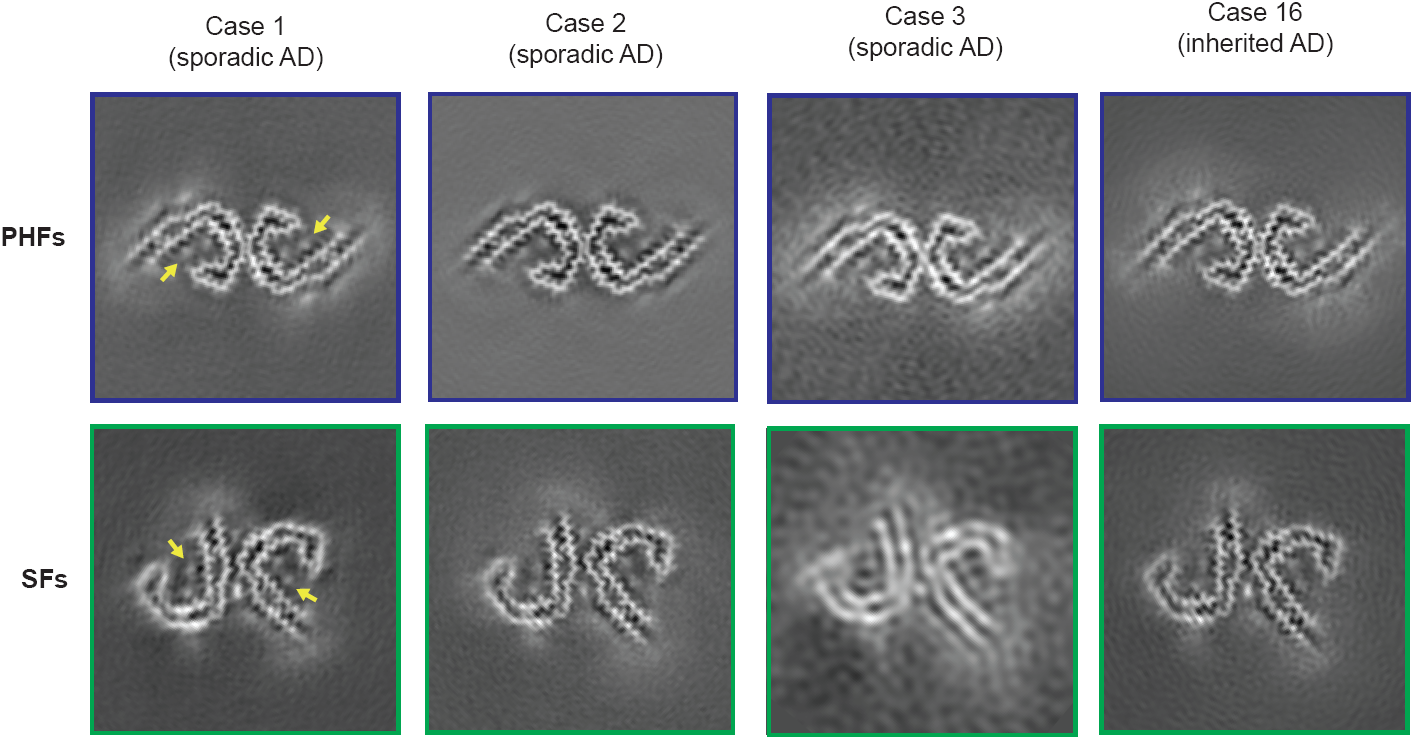
Cryo-EM structures of PHFs and SFs from the frontal cortex of AD cases 1, 2, 3 and 16. All structures show identical pairs of C-shaped protofilaments and the same inter-protofilament packing in PHFs and SFs. Cases 1, 2 and 3 had sporadic AD, whereas case 16 had inherited AD (mutation V717F in *APP*). The filament structures of case 1 are from (19); the structures from cases 2, 3 and 16 are first described here. All cases had a majority of PHFs and a minority of SFs. Yellow arrows indicate the extra densities, which are present in PHFs and SFs from all four cases, bordering the solvent-exposed side chains of R349 and K375, and of H362 and K369.

### Immuno-gold negative-stain electron microscopy

To extend our analysis to a larger number of AD cases and additional brain regions from the first case studied by cryo-EM (19), we used immuno-EM to assess filament morphologies and the labelling of extracted filaments by repeat-specific anti-tau antibodies. We used antibodies specific for R1 (BR136), R2 (Anti-4R), R3 (BR135) and R4 (TauC4) of tau, all of which label tau bands in Western blots of dissociated filaments (19,20) (Figure 3). However, linear epitopes buried in the cores of tau filaments are not accessible to antibodies, in contrast to epitopes located in the fuzzy coat, which give positive labelling (16). Moreover, mild pronase treatment removes the fuzzy coat, abolishing this positive labelling.

**Fig. 3.**
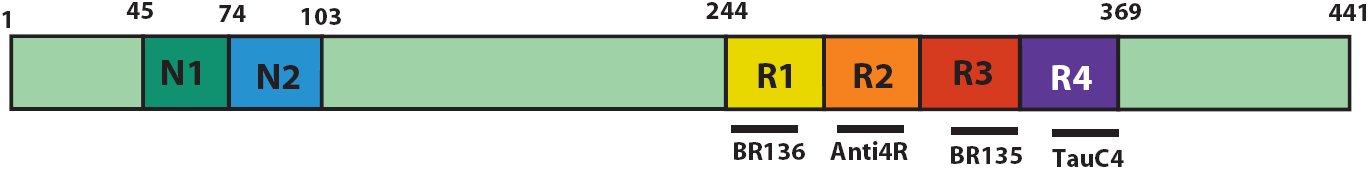
The 441 amino acid isoform of human tau and epitopes of repeat-specific antibodies. The amino-terminal inserts are labelled N1 and N2, with the microtubule-binding repeats being R1-R4. Black lines indicate the epitopes of antibodies BR136 (R1), anti-4R (R2), BR135 (R3) and tauC4 (R4) that were used for immuno-EM.

Besides tau filaments from the four cases analysed by cryo-EM (cases 1-3 and 16), we also looked at tau filaments from the frontal cortex of twelve additional cases of sporadic AD (patients 4-15) and one additional case of dominantly-inherited AD (V717F in *APP*, patient 17) (Table 1). In all cases, extracted filaments consisted of a majority of PHFs and a minority of SFs, as by negative-stain EM. Antibodies BR136 and Anti-4R decorated the filaments from all cases before, but not after, pronase treatment. By contrast, BR135 and TauC4 did not decorate PHFs or SFs from the frontal cortex of cases 4-15 or case 17, either before or after pronase treatment (Figure 4, Online Resource 6). This establishes that R3 and R4 of tau lie in the filament cores. These findings are consistent with the presence of the same tau sequences in the cores of tau filaments in the frontal cortex of all analysed cases (17 cases of sporadic AD and 2 cases of dominantly-inherited AD).

**Fig. 4.**
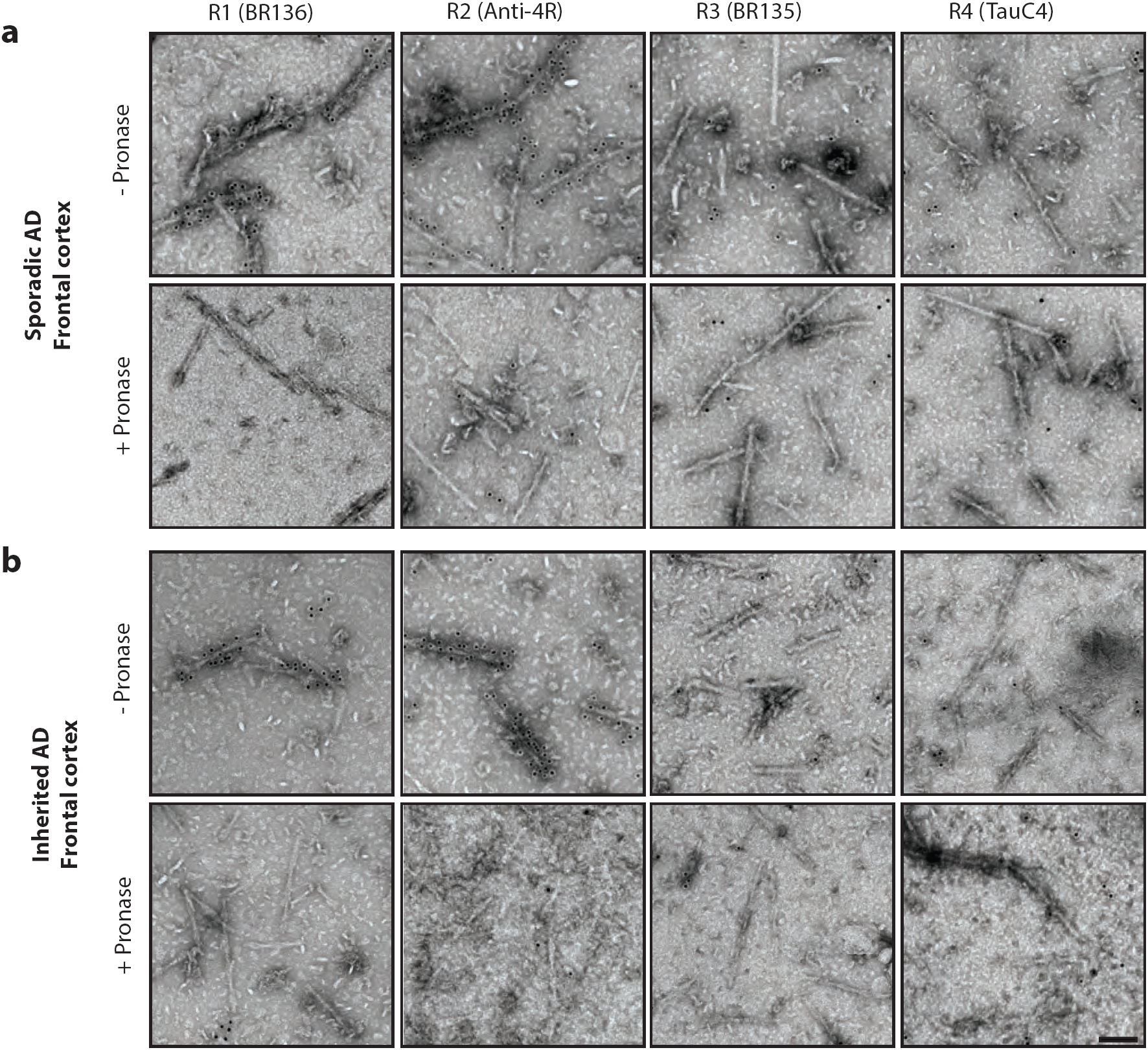
Immuno-EM of tau filaments from the frontal cortex of sporadic and inherited AD cases. Representative images from sporadic (a) and inherited (b) cases of AD before (-) and after (+) pronase treatment. Frontal cortex from fifteen cases of typical sporadic AD, two cases of atypical sporadic AD (PCA) and two cases of typical inherited AD (mutation V717F in *APP*) was studied. Tau filaments from all cases were decorated by BR136 and Anti-4R before pronase treatment, but not by BR135 or TauC4. PHFs were in the majority and SFs in the minority. Scale bar, 100 nm.

To extend our findings to different brain regions, we studied tau filaments from temporal, occipital, parietal and cingulate cortices, as well as from thalamus, substantia innominata and putamen of case 1 of sporadic AD, whose frontal cortex was used for cryo-EM and immuno-EM (Figures 2 and 5, Online Resource 6). In all regions, a majority of PHFs and a minority of SFs were in evidence. By immuno-EM, filaments were labelled by BR136 and Anti-4R before, but not after, pronase treatment, and not by BR135 or TauC4. This establishes that in all brain regions examined, R3 and R4 are present in the core of the filaments.

**Fig. 5.**
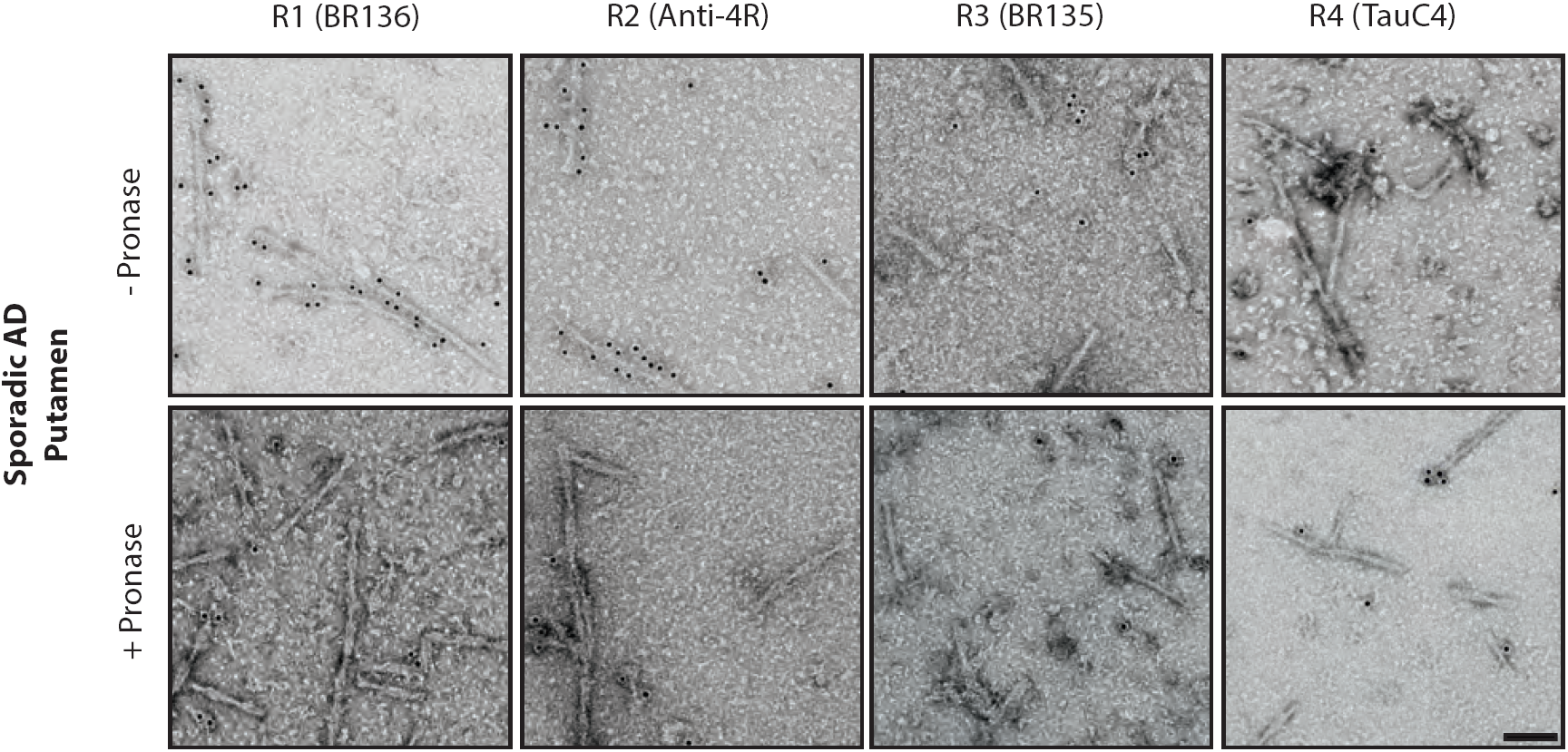
Immuno-EM of tau filaments from the putamen of sporadic AD case 1. Representative images before (-) and after (+) pronase treatment. In addition to frontal cortex and putamen, temporal cortex, parietal cortex, cingulate cortex, thalamus and substantia innominata from case 1 were also analysed. Tau filaments from all brain regions were decorated by BR136 and Anti-4R before pronase treatment, but not by BR135 or TauC4. PHFs were in the majority and SFs in the minority. Scale bar, 100 nm.

PCA is an atypical form of sporadic AD with abundant tau filaments in the occipital cortex (22). As in typical AD, a majority of PHFs and a minority of SFs were present in frontal and occipital cortices of cases 18 and 19. Tau filaments were decorated by BR136 and Anti-4R, but not by BR135 or TauC4 (Figure 6, Online Resource 6). Following pronase treatment, the decoration by BR136 and Anti-4R was lost, establishing also for these cases the presence of R3 and R4 of tau in the filament cores.

**Fig. 6.**
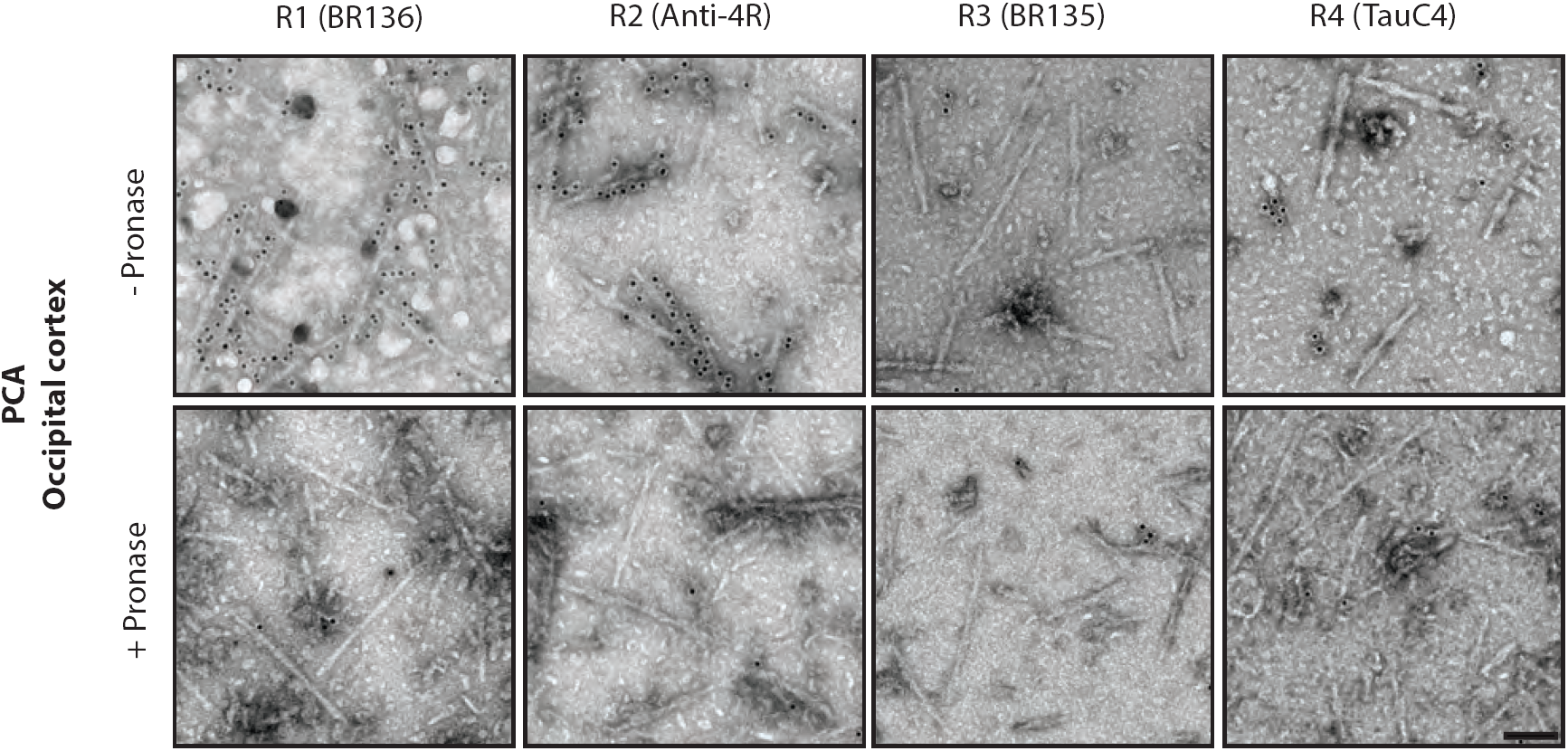
Immuno-EM of tau filaments from the occipital cortex of PCA. Representative images before (-) and after (+) pronase treatment. Occipital cortex from two cases of PCA, an atypical form of sporadic AD, was studied. Case 19 is shown. Tau filaments from both cases were decorated by BR136 and Anti-4R before pronase treatment, but not by BR135 or TauC4. PHFs were in the majority and SFs in the minority. Scale bar, 100 nm.

## DISCUSSION

The cryo-EM structures of the cores of PHFs and SFs from the frontal cortex of four typical cases of AD (three sporadic and one inherited) showed a common protofilament fold. The filament structures from case 1 were reported last year (19). The structures of case 2 were of higher resolution than those of case 1, but the resolutions for cases 3 and 16 were lower. The achievable resolution of filament structures from a given AD case is mainly determined by the numbers of filaments that can be extracted from the brain. PHFs and SFs from all four cases were made of two protofilaments with a common cross-β/β-helix C-shaped architecture. The interfaces between the two protofilaments of PHFs and between those of SFs were the same in all four cases. These findings show that the PHF and SF structures are identical between sporadic (irrespective of *APOE* genotype) and inherited cases of typical AD.

The higher resolution cryo-EM structures of PHFs and SFs from case 2 support the presence of two additional residues at the N-terminus, and two additional residues at the C-terminus of the ordered core of the protofilament. At the C-terminus, these residues were R379 and E380 from the sequence after R4. Because AD filaments contain a mixture of all six brain tau isoforms (16), the additional density at the N-terminus arose from a mixture of 3R and 4R tau isoforms. They comprised G304 and S305 from R2 in 4R tau, and G273 and K274 from R1 in 3R tau. The additional cryo-EM density at the N-terminus was consistent with the presence of a glycine and a mixture of a partially disordered serine and a lysine. As in the structures reported from AD case 1, weaker densities extended from the N- and C- terminal regions of the core, which could accommodate additional tau residues in more dynamic or transiently occupied structures (19). Densities interacting with the sidechains of K317, T319 and K321 were also observed, which we previously suggested could be residues ^7^EFE^9^ of tau forming part of the discontinuous MC1 epitope (19). Similar densities were also found in the structures of tau filaments from PiD (20).

Since cryo-EM is a low-throughput technique, we also used immuno-EM to study tau filaments from the frontal cortex of the four cases used for cryo-EM plus another fifteen cases of AD. They represented both typical and atypical cases, and a variety of *APOE* genotypes. It is important to combine both techniques, because immuno-EM does not provide high-resolution structural information. In all cases, PHFs were in the majority and SFs in the minority. They were decorated by BR136 and Anti-4R, but not by BR135 or TauC4, indicating the presence of R3 and R4 in the filament cores. Importantly, the immuno-EM labelling of tau filaments from the cases used for cryo-EM was indistinguishable from that of the other cases studied, including PCA, an atypical form of sporadic AD. Moreover, for case 1, brain regions other than frontal cortex, including subcortical areas, gave an identical decoration pattern and contained a majority of PHFs. Thus, the cores of all tau filaments contained R3 and R4.

The cryo-EM structures of tau filaments from individuals with end stage AD do not tell us how the assembly process may have started. Since tau is very hydrophilic (24), it is not surprising that the formation of filaments by recombinant protein requires cofactors, such as heparin (25,26). Negatively charged cofactors or chemical modifications of tau may also be needed for the ordered assembly of tau into filaments in human brain. The extra densities bordering the positively charged side chains of solvent-exposed amino acids in the cryo-EM maps of PHFs and SFs from all four cases may be such factors. Once a seed has formed, tau may be able to assemble and propagate (27,28). In a mouse model transgenic for human mutant P301S tau, short tau filaments had the greatest seeding activity in non-neuronal cells expressing full-length human P301S tau (29). It remains to be seen if this is also the case in AD.

The present findings suggest that the filament cores from multiple brain regions of typical sporadic and inherited cases of AD, as well as of atypical sporadic cases, contain identical tau sequences. They are consistent with the existence of a common fold for tau filaments in AD. Tau is an intracellular protein. Its accumulation may involve mechanisms that differ from those that facilitate the seeding of proteins that form amyloids extracellularly. For PrP^Sc^ and Aβ, structurally heterogeneous assemblies, also referred to as conformational clouds, have been described (30-32). Knowledge of the structural differences between assemblies of PrP^Sc^ and Aβ must await the determination of their high-resolution structures from brain.

Distinct tau filament folds are found in specific diseases. The cryo-EM structures of tau filaments from the brain of an individual with PiD revealed that they adopt a fold that is different from the AD fold (20). Immuno-EM results for eight additional cases of PiD revealed a common ordered filament core comprising R1, R3 and R4. Therefore, specific molecular conformers of aggregated tau may define different diseases, without there being significant variation in filament structures between individuals with a given disease. This understanding will inform the design of diagnostic ligands of higher specificity and the development of therapies targeting tau assembly. It remains to be seen if what is true of AD and PiD also applies to other human tauopathies.

## MATERIALS AND METHODS

### Extraction of tau filaments

Sporadic cases of typical AD were selected according to several criteria, including a wide spread in the ages at death of patients and in the post mortem intervals (time between death and brain autopsy), a tau pathology of Braak stage 6 and no or only mild co-pathologies. Inherited cases of typical AD were chosen based on the presence of a V717F mutation in *APP* and sporadic cases of atypical AD exhibited a clinical and neuropathological picture of PCA. Neurohistology and immunohistochemistry were carried out as described (33).

Sarkosyl-insoluble material was extracted according to (16). Approximately 6 g frontal cortex was used for cryo-EM and 0.6 g from each brain region studied for immuno-EM. The pelleted sarkosyl-insoluble material was resuspended in 10 mM Tris-HCl pH 7.4, 800 mM NaCl, 5 mM EDTA, 1 mM EGTA, with a final concentration of 10% (w/v) sucrose at 750 ml per g tissue, followed by centrifugation at 20,100 g for 30 min at 4^°^ C. The pellet was resuspended in 20 mM Tris-HCl pH 7.4 containing 100 mM NaCl at 250 ml/g tissue and centrifuged at 100,000 g for 30 min at 4^°^ C. The final pellet was resuspended in buffer at 15 ml/g tissue for cryo-EM and 150 ml/g tissue for immuno-EM.

### Electron cryo-microscopy

Extracted tau filaments were applied to glow-discharged holey carbon grids (Quantifoil Au R1.2/1.3, 300 mesh) and plunge-frozen in liquid ethane using an FEI Vitrobot Mark IV. Images were acquired on a Gatan K2-Summit detector in counting mode using an FEI Titan Krios at 300 kV. A GIF-quantum energy filter (Gatan) was used with a slit width of 20 eV to remove inelastically scattered electrons. Further details are given in Online Resource 2.

### Helical reconstruction

Movie frames were corrected for gain reference, motion-corrected and dose-weighted using MOTIONCOR2 (34). Aligned, non-dose-weighted micrographs were used to estimate the contrast transfer function in Gctf (35). All subsequent image-processing steps were performed using helical reconstruction methods in RELION 2.1 (23,36). Filaments were picked manually. For case 2, PHFs and SFs were picked as separate datasets. For cases 3 and 16, filaments were collected as a single dataset; PHFs and SFs were subsequently separated by reference-free 2D classification of segments comprising an entire helical crossover. PHF and SF segments were re-extracted using box sizes of 200 or 270 pixels and an inter-box distance of approximately 14 Å. Reference-free 2D classification was performed and segments contributing to suboptimal 2D class averages were discarded. For cases 2 and 16, initial 3D models were constructed *de novo* from 2D class averages of segments comprising entire helical crossovers, low-pass filtered to 40 Å. For case 3, the PHF and SF reconstructions from case 1 (19) were used as initial 3D models, low-pass filtered to 10 Å. 3D classification with local optimisation of helical twist and rise was performed to remove segments contributing to suboptimal 3D class averages, and the selected segments were used for 3D auto-refinement with optimisation of the helical twist. A value of 10% was used for the helical z percentage parameter. For case 16, particle polishing was performed, followed by further 3D auto-refinement. The final reconstructions were sharpened using standard post-processing procedures in RELION and helical symmetry was imposed using the RELION helix toolbox (23). Final, overall resolution estimates were calculated from Fourier shell correlations at 0.143 between two independently refined half-maps, using phase randomization to correct for convolution effects of a generous, soft-edged solvent mask (37). Local resolution estimates were obtained using the same phase randomization procedure, but with a soft spherical mask that was moved over the entire map. For further details, see Online Resource 2.

### Model building and refinement

Compared to (19), the PHF and SF reconstructions from the frontal cortex of AD patient 2 showed clear densities corresponding to two additional residues at the N-terminus (G273 and K274 from 3R tau; or G304 and S305 from 4R tau) and two residues at the C-terminus (R379 and E380) of the protofilament core. These residues were added to the PHF and SF models from the frontal cortex of AD patient 1 (19) (PDB accession numbers 503L and 503T) using COOT (38). The new models were then refined against the PHF and SF reconstructions from patient 2 using targeted real-space refinement in COOT. The models were subsequently translated to give stacks of three consecutive monomers in order to preserve nearest-neighbour interactions for the middle chains in subsequent Fourier space refinements in REFMAC (39). Local symmetry restraints were imposed to keep all β-strand rungs identical. Side-chain clashes were detected using MOLPROBITY (40) and corrected by iterative cycles of real-space refinements in COOT and Fourier space refinements in REFMAC. Separate model refinements were performed against single half-maps, and the resulting models compared with the other half-maps to confirm the absence of overfitting. The final models were stable in refinements without additional restraints. Further details are given in Online Resource 3.

### Immuno-gold negative-stain electron microscopy

Extracted tau filaments were deposited on glow-discharged 400 mesh formvar/carbon film-coated copper grids (EM Sciences CF400-Cu) for 40 s, blocked for 10 min with PBS + 0.1% gelatin and incubated with primary antibody (1:50) in blocking buffer, essentially as described (16). Primary antibodies were BR136 (raised against residues 244-257) (20), Anti-4R (raised against residues 275-291, with D279) (41), BR135 (raised against residues 323-335) (42) and TauC4 (raised against residues 354-369) (43). Where stated, grids were incubated for 5 min with 0.4 mg/ml pronase (Sigma) in PBS at room temperature and washed, prior to blocking. Following incubation with primary antibodies, they were washed with blocking buffer and incubated with 10 nm gold-conjugated anti-rabbit IgG (Sigma) diluted 1:20 in blocking buffer. The grids were then washed with water and stained with 2% uranyl acetate for 40 s. Images were acquired at 11,000x and 15,000x, with a defocus value of -1.4 mm with Gatan Orius SC200B or Gatan Ultrascan 1,000 CP CCD detectors using a Tecnai G2 Spirit at 120 kV.

### Whole exome sequencing

Genomic DNAs of the AD cases were sequenced at the Center for Medical Genomics of Indiana University School of Medicine. Target enrichment made use of the SureSelectXT human all exon library (V6, 58Mb, Agilent) and high-throughput sequencing using a HiSeq4000 (2x75bp paired-end configuration, Illumina). Bioinformatics analyses were performed as described (44). There were no pathogenic mutations in *MAPT*. Table 1 summarises the findings for *APP*, *PSEN1*, *PSEN2* and *APOE*.

## AUTHOR CONTRIBUTIONS

B.G. performed neuropathology; H.J.G. and R.V. carried out genetic analysis; B.F. and M.S. extracted tau filaments; B.F. and W.Z. performed cryo-EM; B.F., W.Z., A.G.M. and S.H.W.S. analysed cryo-EM data; B.F. refined atomic models; B.F., W.Z. and M.S. conducted immuno-EM; S.H.W.S. and M.G. supervised the project.

## ACKNOWLEDGEMENTS

We thank the patients’ families for donating brain tissues; M.R. Farlow for clinical evaluation; F. Epperson, R.M. Richardson and U. Kuederli for brain collection and analysis; A.W.P. Fitzpatrick for cryo-EM data collection of AD cases 2 and 3; M. Hasegawa for antibody TauC4; S. Chen, C. Savva, G. Cannone and K. Dent for support with electron microscopy; T. Darling and J. Grimmett for help with computing; R.A. Crowther for helpful discussions. M.G. is an Honorary Professor in the Department of Clinical Neurosciences of the University of Cambridge. This work was supported by the UK Medical Research Council (MC_UP_A025_1013 to S.H.W.S. and MC_U105184291 to M.G.), the European Union (Joint Programme-Neurodegeneration Research REfrAME to B.F. and M.G. and the Innovative Medicines Initiative 2 IMPRiND, project 115881, to M.G.), Eli Lilly and Company (to M.S. and M.G.), the US National Institutes of Health (grant P30-AG010133 to B.G.), the Department of Pathology and Laboratory Medicine, Indiana University School of Medicine (to B.G.) and an Alzheimer’s Association Zenith Award (to R.V.). We acknowledge DIAMOND for access to and support of the cryo-EM facilities at the UK electron Bio-Imaging Centre (eBIC), proposal EM17434, funded by the Wellcome Trust, the MRC and the BBSRC.

---

**Online Resource 1.**
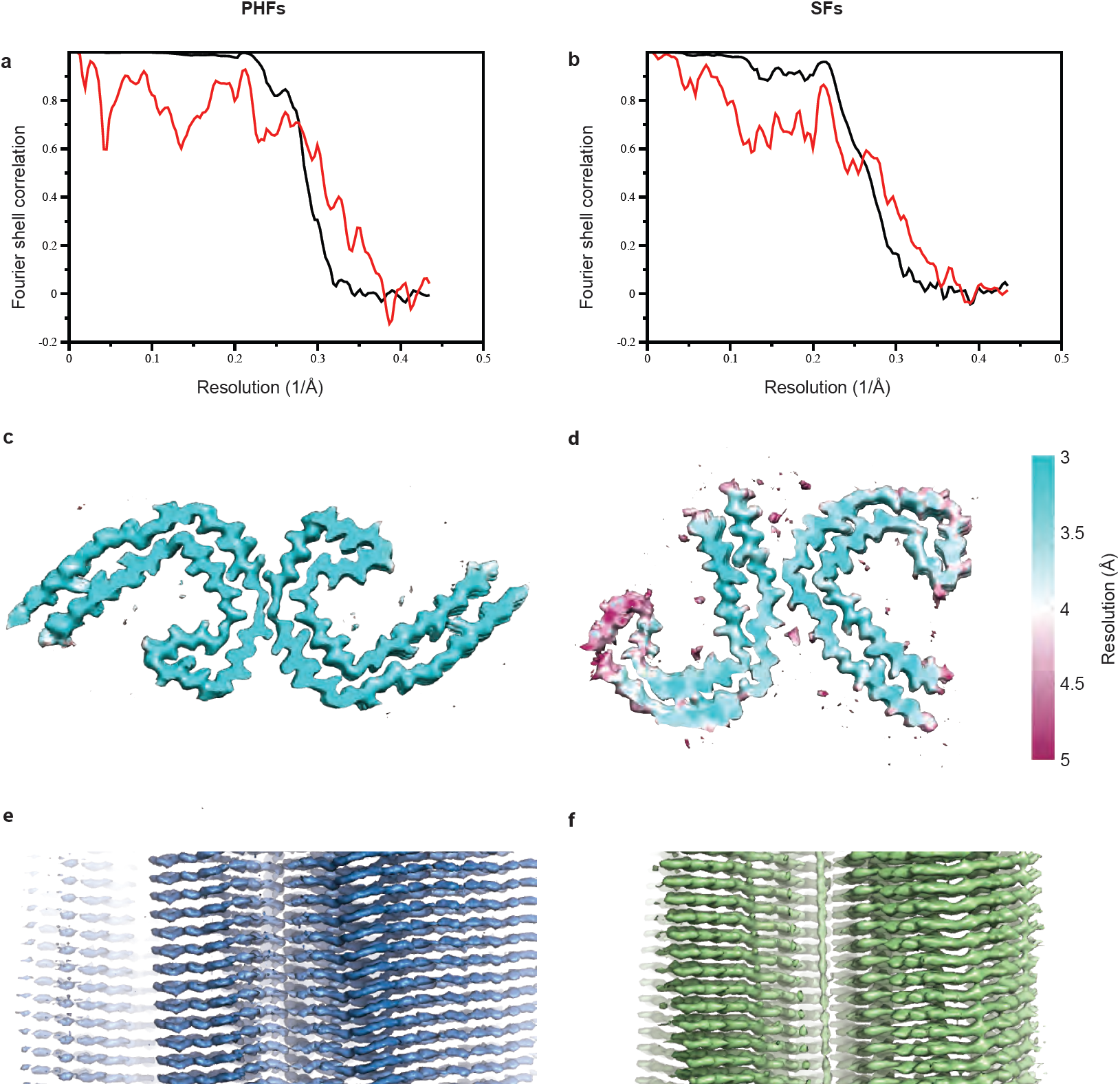
PHFs and SFs from the frontal cortex of AD case 2. (a,b) Fourier shell correlation curves between two independently refined half-maps (black line) and between the cryo-EM reconstruction and refined atomic model (red line) for PHFs (a) and SFs (b). (c,d) Local resolution estimates for PHF (c) and SF (d) reconstructions. (e,f) Views normal to the helical axis of PHF (e) and SF (f) reconstructions.

**Online Resource 2:** Electron cryo-microscopy structure determination

|  | Case 2 |  | Case 3 |  | Case 16 |  |
| --- | --- | --- | --- | --- | --- | --- |
| Data Collection | PHFs | SFs | PHFs | SFs | PHFs | SFs |
| Magnification | ×105,000 | ×105,000 | ×105,000 | ×105,000 | ×165,000 | ×165,000 |
| Defocus range (μm) | -1.7 to -2.8 | -1.7 to -2.8 | -1.7 to -2.8 | -1.7 to -2.8 | -1.7 to -2.8 | -1.7 to -2.8 |
| Voltage (kV) | 300 | 300 | 300 | 300 | 300 | 300 |
| Microscope | Titan Krios | Titan Krios | Titan Krios | Titan Krios | Titan Krios | Titan Krios |
| Detector | K2 Summit | K2 Summit | K2 Summit | K2 Summit | K2 Summit | K2 Summit |
| Frame exposure time (ms) | 200 | 200 | 200 | 200 | 250 | 250 |
| Number of frames | 50 | 50 | 50 | 50 | 44 | 44 |
| Total dose (e<sup>-</sup>/Å<sup>-2</sup>) | 60 | 60 | 60 | 60 | 57.5 | 57.5 |
| Pixel size (Å) | 1.15 | 1.15 | 1.15 | 1.15 | 0.85* | 0.85* |
| Reconstruction |  |  |  |  |  |  |
| Box size (pixel) | 270 | 270 | 270 | 270 | 200 | 200 |
| Inter-box distance (Å) | 28 | 28 | 28 | 28 | 20 | 20 |
| Segments extracted | 197,500 | 62,782 | 35,400 | 35,400 | 190,409 | 190,409 |
| Segments after Class2D | 95,917 | 29,536 | 22,150 | 12,690 | 115,669 | 47,437 |
| Segments after Class3D | n/a | 27,103 | n/a | n/a | 54,900 | 29,594 |
| Resolution (Å) | 3.2 | 3.3 | 4.7 | 8.4 | 3.8 | 4.2 |
| B-factor (Å<sup>2</sup>) | -90 | -61 | -152 | n/a | -87 | -58 |
| Helical rise (Å) | 2.37 | 4.76 | 2.36 | 4.74 | 2.39 | 4.84 |
| Helical twist (°) | 179.45 | -1.04 | 179.40 | -1.05 | 179.45 | -1.02 |

\* For image processing, the segments for case 16 were downscaled to a pixel size of 1.53 Å.

**Online Resource 3:** Patient 2 PHF and SF model statistics

| Refinement | PHFs | SFs |
| --- | --- | --- |
| Initial model used (PDB code) | 5o3l | 5o3t |
| Model resolution (Å) | 3.2 | 3.3 |
| FSC threshold | 0.5 | 0.5 |
| Map sharpening *B* factor (Å<sup>2</sup>) | -90 | -61 |
| Model composition |  |  |
| Non-hydrogen atoms | 587 | 587 |
| Protein residues | 456 | 456 |
| Average *B* factor (Å<sup>2</sup>) | 72.8 | 82.0 |
| R.m.s. deviations |  |  |
| Bond lengths (Å) | 0.019 | 0.016 |
| Bond angles (°) | 1.37 | 1.11 |
| Validation |  |  |
| MolProbity score | 1.90 | 2.00 |
| Clashscore | 5.59 | 6.71 |
| Poor rotamers (%) | 0 | 0 |
| Ramachandran plot |  |  |
| Favored (%) | 88 | 87 |
| Allowed (%) | 100 | 100 |
| Disallowed (%) | 0 | 0 |

**Online Resource 4.**
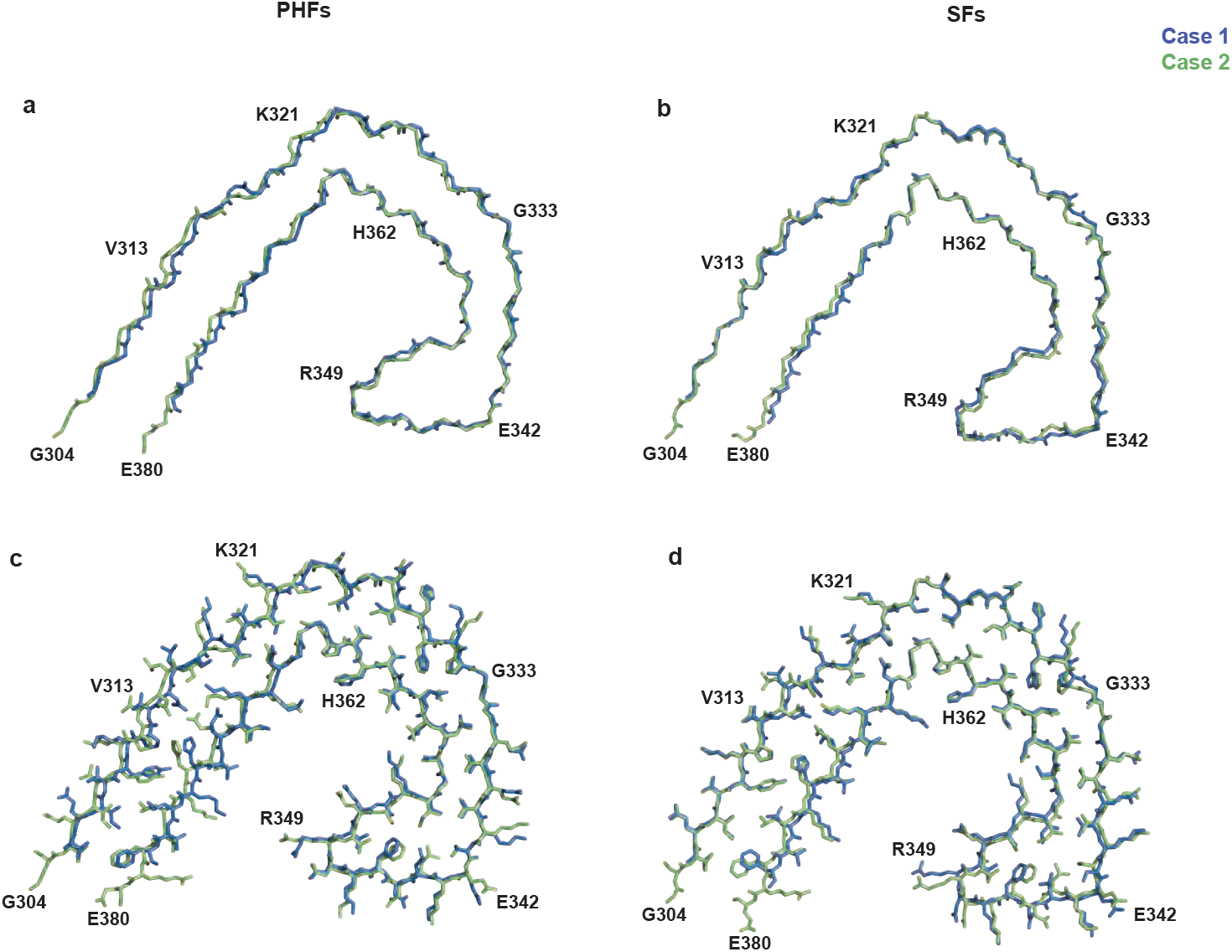
Protofilament structures of AD cases 1 and 2. (a,b) Overlay of backbone atoms of the protofilament structures of PHFs (a) and SFs (b) from AD cases 1 (blue) and 2 (green). (c,d) As in (a,b), but showing all atoms.

**Online Resource 5.**
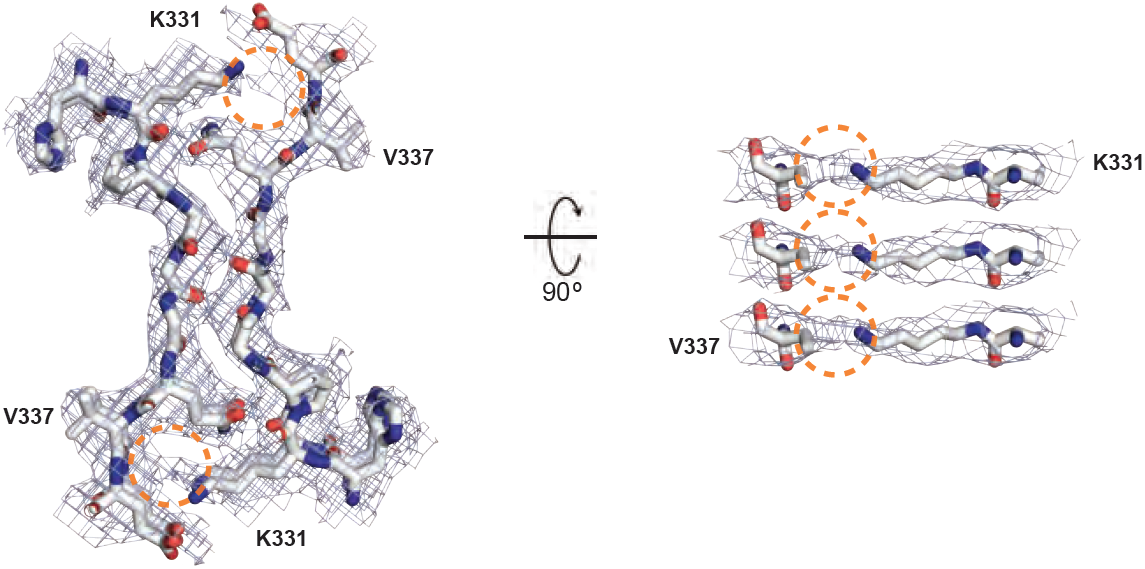
Protofilament interface in PHFs. Packing between residues ^331^KPGGGQV^337^ of the two protofilaments. Inter-protofilament hydrogen bonds are present between G333 and G334, and between Q336 and the backbone carbonyl of K331. Additional densities between the side chains of K331 and the backbone atoms of V337 are highlighted with dashed orange outlines.

**Online Resource 6:**
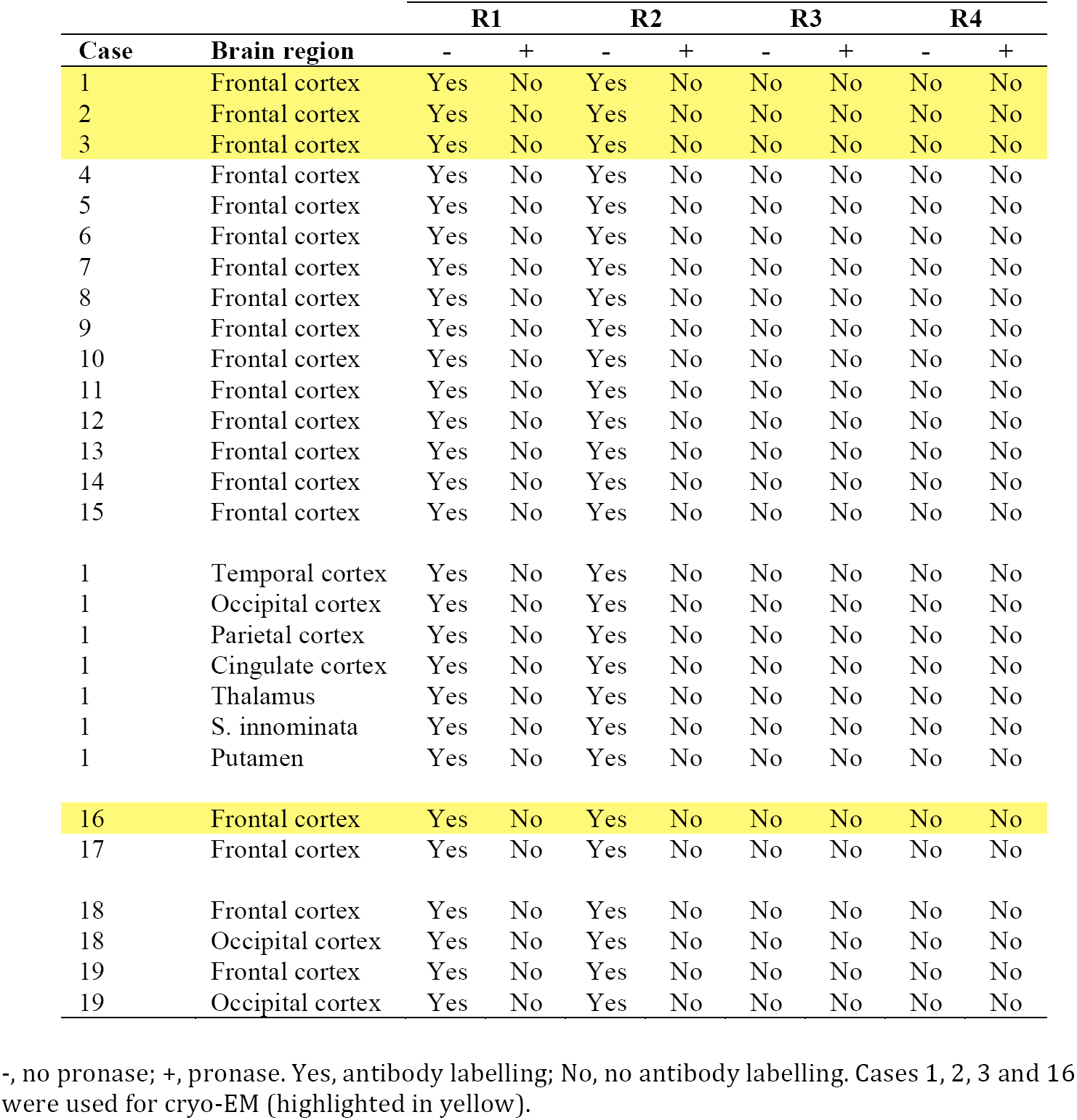
Immuno-EM labelling of PHFs and SFs

